# High fidelity genetic markers for sexing *Cannabis sativa* seedlings

**DOI:** 10.1101/2024.09.10.612257

**Authors:** Djivan Prentout, Salma El Aoudati, Fabienne Mathis, Gabriel AB Marais, Hélène Henri

## Abstract

The uses of *Cannabis sativa*, a dioecious species with an XY sex chromosome system, are varying from fiber and oil to cannabinoids, among others. In most cases, males are undesirable and the sexual dimorphism at immature plants is too subtle for reliable phenotypic sexing, making genetic approaches promising. In this technical note, we present a multiplex PCR-mix, that includes two markers of Y-specific coding regions and one autosomal control marker. This PCR-mix, tested across 12 hemp-type cultivars, encompassing approximately 200 individuals, achieved a 99.5% success rate in identifying the sex of *C. sativa* seedlings.

## Introduction

*Cannabis sativa* is one of the oldest domesticated plants and is used in a wide range of purposes. Two primary groups of cultivars have been described: hemp-types and drug-types (Ren *et al*., 2021; Rehman *et al*., 2021). Hemp-type cultivars are typically used for textiles, food or oilseed production, whereas drug-types are cultivated for medicinal and recreational uses (Rehman *et al*., 2021). The recent global legalization of the latter applications has led to a significant increase of the drug-type production in the past decade. As a consequence, the global economy of *C. sativa* now surpasses tens of billions of dollars annually (Malabadi *et al*., 2024).

In nature, *C. sativa* is dioecious (*i.e*. male and female flowers are on separate individuals) and the sex is determined by an XY pair of sex chromosomes (Ming *et al*., 2011; Divashuk *et al*., 2014). Female plants are preferred for most of the purposes listed before. For example, female plants produce thicker fibers and are therefore favored for textile production. More critically, males do not produce cannabinoids (THC and CBD) in high concentration, and in case of a pollination, females cease their production (Chandra *et al*., 2017). This motivates an early identification of males in crops, and more generally, a better control of the sex in *C. sativa* (Chandra *et al*., 2017).

Three primary strategies can be employed to remove male plants from a crop (i) sex reversal of the individuals, (ii) vegetative propagation (i.e. cloning) of females, or (iii) identification, and withdrawing, of male individuals from the crops. Sex reversal typically implies the administration of a silver agent to females, in order to produce hermaphrodite flowers (see Chandra *et al*., 2017). Such hermaphrodites have two X chromosomes, leading their self pollination to produce only “feminized” seeds (with a success rate of 95%, see Chandra *et al*., 2017). However, besides being pricey this process is unstable throughout generations (reviewed in (Chandra *et al*., 2017; Malabadi *et al*., 2023)). Vegetative propagation of female individuals requires highly controlled conditions to avoid sex plasticity. Additionally, somatic mutations may accumulate over generations, decreasing disease resistance or cannabinoid content(Trancoso *et al*., 2022). Phenotypic identification of the sex in immature *C. sativa* plants is nearly impossible due to the lack of sexual dimorphism in young individuals. Genetic approaches offer a promising alternative. Recently, a sex marker utilizing a PACE-PCR Allele competitive Extension showed a high success in sex identification (Toth *et al*., 2020). However, this marker, along with previous markers, is located in retrotransposons, which may affect its accuracy across different cultivars (Törjék *et al*., 2002; Sakamoto *et al*., 2005; Toth *et al*., 2020).

In a previous analysis, we conducted a transcriptome-wide segregation analysis of *C. sativa* to identify genes located in the non-recombining region of the sex chromosomes (hereafter referred to as sex-linked genes; Prentout *et al*., 2020). We identified more than 550 sex-linked genes and showed that some regions of the sex chromosomes are highly divergent (X-Y synonymous divergence reaches 40%)(Prentout *et al*., 2020). These highly divergent regions are of particular interest because they could be shared in all *C. sativa* populations/cultivars, providing a unique opportunity to develop universal Y-linked markers for an early sex identification. In the present study, we use these divergent sex-linked genes to develop a multiplex PCR assay aimed at identifying males in *C. sativa* seedlings.

## Material and Methods

### Design and *in silico* validation of the primers

Among the sex-linked genes identified in (Prentout *et al*., 2020), we selected the XY gene pairs with high synonymous divergence and a length greater than 300 pb. For each gene pair, we aligned the Y-linked sequence from the cross we generated (cultivar Zenitsa, hemp-type, see Prentout *et al*., 2020) with three X-linked sequences: (i) one from the same cross as the Y sequence, (ii) one from a Purple Kush cultivar (drug-type) and (iii) one from a Finola cultivar (hemp-type) (the two latter from van Bakel *et al*., 2011). This increased the probability of identifying X-Y divergent sites shared by different populations. We aligned the sequences with the Qiagen CLC Main workbench tool ‘create alignment’ (version 8.0.1, see https://www.qiagenbioinformatics.com) and used the tool ‘Design Primer’ (Krämer *et al*., 2014). We kept the coding regions with at least two fixed differences between X-linked and Y-linked sequences, and avoided complementary forward and reverse primers to avoid association between our primers during PCR.

We used the *in silico* PCR tool from the van Bakel lab to test the size of the amplicons, and to verify that a unique region of the genome was amplified (http://genome.ccbr.utoronto.ca/). Our primers were tested *in silico* on both the Purple Kush and the Finola genomes. As both genomes are female genomes, we expected an amplification of X-linked primers but not of the Y-linked.

### Plant material for *in vitro* validation

We collected 192 samples from twelve hemp-type cultivars. The sampling included both accessions and varieties. Seeds were sown in a climatic chamber and exposed to 18 hours of light and 6 hours of darkness, at 22^°^C and 18^°^C, respectively. Flowering was induced by altering the light cycle to 12 hours of daylight, with the same temperature conditions as the initial cycle. Once the plants could be sexed phenotypically, we collected eight leaf discs per sample and air-dried them for DNA isolation.

### DNA extraction

We used the CTAB-method from Koh *et al*., 2021 protocol (Koh *et al*., 2021). From 10 to 30mg of leaves were placed in a 2mL round bottom tube, with a 5mm stainless steel bead, and placed for at least 15 minutes at -80^°^C. Frozen samples were ground into powder by setting the Tissue Lyser (Retsch) speed at 20Hz for 2 minutes. 400µL of CTAB DNA extraction buffer (2% w/v CTAB, 20 mM EDTA.Na_2_.2H_2_O, 1,4 M sodium chloride and 100 mM tris) was then added to the tube supplemented with 0.3% (v/v) β-mercaptoethanol. Tubes were incubated at 65^°^C for 30 minutes in a thermomixer (Eppendorf) under agitation at 400rpm. Debris was separated from the liquid part through centrifugation (15000 × g for 5 min at RT). The resulting supernatant (270µL) was transferred to a new 1.5 mL tube, and an equal volume of 24:1 chloroform:isoamyl alcohol was added. The tubes were constantly inverted for 5 min to mix the solution, and the aqueous phase was separated by centrifugation (15000 × g for 5 min at 4^°^C). 240µL of the recovered aqueous phase was transferred to a new 1.5 mL tube and RNA was digested with the addition of 0.5 μL of 10 mg/mL of RNAseA and incubated for 15 minutes at 37^°^C. Chloroform:isoamyl alcohol extraction was repeated, and 240μL of the resulting supernatant was transferred to a new tube. DNA was precipitated by the addition of ½ volume (120 μL) of 5 M NaCl and three volumes (720μL) of ice-cold absolute ethanol and incubation at -20^°^C for at least 30 min. Tubes were centrifuged at 15000 × g for 5 min at 4^°^C to pellet the DNA precipitates. Pellets were washed twice with 500 μL of 70% ethanol (centrifuged at 15000 × g for 5 min at 4^°^C) and resuspended in 50 μL of nuclease-free water.

### PCR reaction

The DNA extracts were diluted 1:10 before being used for the PCR reactions. PCR reactions were performed in a final volume of 10µL using 5µL of Taq Ozyme HS mix 2X (Ozyme, ref #OZYA006), 1.5µL of primer mix comprising 10µM of each primer, 1.5µL of ultrapure water, and 2µL of 1:10 diluted DNA. The cycling conditions consisted of an initial denaturation step lasting 1 min at 95^°^C, followed by 35 cycles of denaturation at 95^°^C for 15 s, annealing at 58^°^C for 15 s, and elongation at 72^°^C for 30 s, with a final extension step at 72^°^C for 5 min.

## Results

We selected 2 Y-linked primers and 1 autosomal primer in order to develop a multiplex approach (Table 1). The autosomal primer is used to control if the PCR worked, and the two Y-linked primers increase the success probability for distant cultivars. This protocol was tested on 192 samples (96 males and 96 females) from 12 different hemp cultivars (Table 2). It is worth noting that the number of samples tested for each cultivar is ranging between 1 and 33 (Table 2). Among the 192 tested samples, the sex was correctly inferred for 191 of them, meaning a success rate of 99.5%. Only one female individual was detected as a male (red cell in Table 2).

**Table 1.**
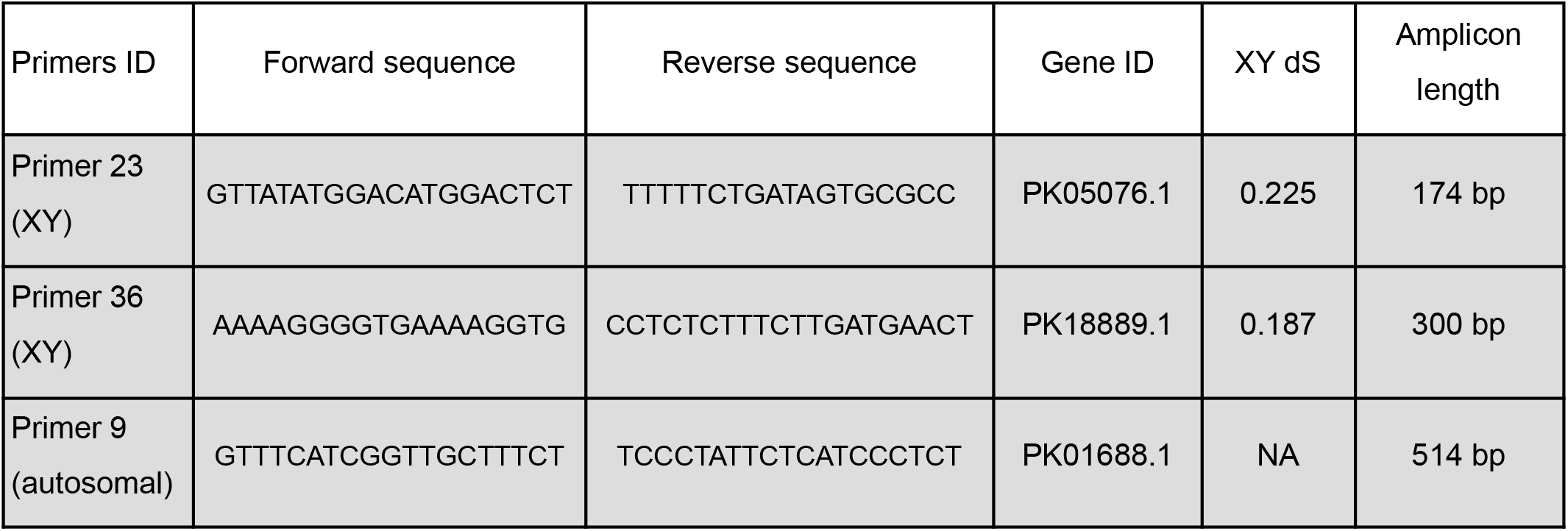
Primer IDs and sequences along with the gene ID they belong to, the segregation type, the XY divergence and the amplicon length.

**Table 2.** Number of samples tested for each of the twelve cultivars. The table reports: the number of samples tested, the number of individuals correctly sexed for males, the number of individuals correctly sexed for females, the number of false negatives (male sexed as female), the number of false positives (female sexed as male), the number of cases for which the PCR failed, and the success rate.

| Cultivar | Number of sample | Male sexed as male | Female sexed as female | Male sexed as female | Female sexed as male | Experiment failed | Success rate (%) |
| --- | --- | --- | --- | --- | --- | --- | --- |
| Pu | 22 | 10 | 12 | 0 | 0 | 0 | 100 |
| Cr | 30 | 15 | 14 | 0 | 1 | 0 | 97 |
| Ch | 8 | 4 | 4 | 0 | 0 | 0 | 100 |
| D1 | 33 | 15 | 18 | 0 | 0 | 0 | 100 |
| Ti | 18 | 12 | 6 | 0 | 0 | 0 | 100 |
| Ma | 12 | 6 | 6 | 0 | 0 | 0 | 100 |
| Ks | 29 | 15 | 14 | 0 | 0 | 0 | 100 |
| D8 | 29 | 15 | 14 | 0 | 0 | 0 | 100 |
| D2 | 1 | 0 | 1 | 0 | 0 | 0 | 100 |
| Ca | 1 | 0 | 1 | 0 | 0 | 0 | 100 |
| No | 1 | 0 | 1 | 0 | 0 | 0 | 100 |
| Fi | 8 | 4 | 4 | 0 | 0 | 0 | 100 |
| Total | 192 | 96 | 95 | 0 | 1 | 0 | 99.5 |

The primers have been tested only with hemp-type cultivars. We explored whether the PCR multiplex mix could be used in drug-type cultivars of *C. sativa*. To do so, we ran an *in silico* amplification using the *Primer-BLAST tool* in the nr database (Sayers *et al*., 2021) (see https://www.ncbi.nlm.nih.gov/tools/primer-blast/index.cgi). We selected the genomes for which the sex was reported, and we used the default parameters for the amplification. A total of eight female genomes and three male genomes were available for *C. sativa* (see Supplementary Table 1). The primers for autosomal locus #9, our internal control, successfully produced the expected size amplification for all genomes. None of the female genomes resulted in the amplification of any Y-specific primers. The sex of the three male genomes (including two a drug-type and one hemp-type) were correctly identified by primer #23. However, the primer #36, successfully amplified only in the hemp-type genome. This suggests that at least one of the two Y-specific primers could be effective in identifying the sex in drug-type cultivars of *C. sativa*.

## Discussion

In the present study, we developed a PCR multiplex mix, which includes two Y-specific markers and one autosomal marker to identify the sex of *Cannabis sativa* samples. The multiplex mix correctly identifies the sex in 99.5% of the cases, which makes it among the most accurate methods of the Cannabaceae family (Toth *et al*., 2020; Clare *et al*., 2024). Only one female sample was identified as a male. Given that the sex was correctly assigned for 15 males and 14 females of the same cultivar, we cannot rule out the hypothesis of an error in the phenotypic identification of the sex, or maybe a sex reversion of the individual (genotypic male with a female phenotype). Furthermore, our method offers a 100% specificity (i.e. absence of males assigned as females), which is crucial to avoid males in crops.

Although we only tested the mix in hemp-type cultivars, the *in silico* analysis showed that one of the two Y-specific markers amplified in drug-type genomes obtained from male individuals. The *in silico* analysis relies on the quality of the Y assemblies and it is possible that the other Y-specific marker did not amplify because of assembly errors. This suggests that the mix could be efficient to identify sex of most, if not all, *C. sativa* cultivars. Future work should aim at testing our markers in drug-type cultivars.

## Acknowledgement

We thank Genady Karlov and his team, Jernej Jakse and Marko Flajšman for discussions and for some preliminary tests not shown in this manuscript.

## Author contribution

Conceptualization: G.A.B.M, H.H

Methodology: D.P, G.A.B.M, H.H

Software: D.P, H.H

Formal analysis: D.P, S.E.A, H.H

Investigation: D.P, S.E.A, F.M, G.A.B.M, H.H

Data curation: D.P, S.E.A, H.H

Ressources: F.M

Writing original draft: D.P, G.A.B.M, H.H

Review and editing: All authors

